# Influence Of Desire To Belong And Feelings Of Loneliness On Emotional Prosody Perception In Schizophrenia

**DOI:** 10.1101/092080

**Authors:** Rachel Mitchell, Krystal Gamez, Christian Kohler, Monica E. Calkins, Bruce I. Turetsky, David I. Leitman

**Author notes:** Address correspondence to: David I. Leitman, Ph.D. Department of Psychiatry Neuropsychiatry Program - Brain Behavior Laboratory University of Pennsylvania Gates Pavilion 10th floor 3400 Spruce St Philadelphia, PA 19104-4283 E. The Tower of Babel (Pieter Bruegel the Elder (1563) - illustrating human collaboration) (David I. Leitman).

## Abstract

<u>Objective</u>: Humans are social creatures, with desires to connect or belong, producing loneliness when isolated. Individuals with schizophrenia are often more isolated than healthy adults and demonstrate profound social communication impairments such as vocal affect perception (prosody). Loneliness, levels of desire for social connectedness (need to belong, NTB), and their relationship to perception of social communications have not been investigated in schizophrenia.

<u>Method</u>: In a sample of 69 individuals (36 SZ), we measured endorsements of loneliness and NTB, and evaluated their putative relationships to clinical symptoms and social communication abilities, as indexed by emotional prosody and pitch perception.

<u>Results</u>: Loneliness endorsement was highly variable but particularly so in patients, whilst patients endorsed NTB at levels equivalent to healthy controls. In schizophrenia, pitch and prosody acuity were reduced, and prosody perception correlated with NTB. Loneliness, but not desire for social connectedness, correlated with negative symptoms.

<u>Conclusion</u>: Loneliness and negative symptoms likely exert bidirectional effects on each other. Loneliness and desire to form interpersonal attachments may be pivotal in shaping and stimulating social interactions and, subsequently, the ability to perceive social intent through prosody. Intact NTB levels in patients augurs well for cognitive remediation which, target vocal-communication processing to improve social skills.

**Significant Outcomes:**

1. Patients with schizophrenia endorsed higher levels of loneliness than controls, but ratings of desire for social connectedness were at normal levels.
2. Pitch acuity and prosody perception were correlated, confirming the importance of basic sensory processing in recognizing prosodic emotions.
3. Socio-cognitive perceptual ability (emotional prosody perception) correlated with increased desire for social connections, implying that they may still be motivated to find social interactions reinforcing. Thus interventions to improve perceptual deficits could still be an effective means of improving social function.

**Limitations:**

1. Causal relationships between desire for social connections, loneliness, and emotional prosody perception cannot be inferred through correlations and cross-sectional studies alone.
2. Subjective endorsements of loneliness through self-report are not the same thing as objective indices of loneliness. New and more extensive tools for measuring desire for both loneliness and social connectedness may be needed.
3. Direct experimental comparison of the interrelations between desire for social connectedness, loneliness, pitch acuity and emotional prosody perception in patients with schizophrenia and other populations such as autism will enable a more accurate comparison of the likely success of remediating socio-cognitive perceptual impairment in neuropsychiatric disorders.

## 1. INTRODUCTION

A singular aspect of human evolutionary fitness is our ability to collaborate with each other, and it is this innate drive to collaborate that sparked our unique ability to communicate through language (1). From such a perspective, it is not surprising that the social impairments in neuropsychiatric illnesses such as autism and schizophrenia (SZ) often foreshadow poor functional outcome(2–7).

A crucial element of social interaction is the ability to perceive the intentions of others. The facial, prosodic, and gestural expressions of others serve as communicative signals, indicating a person’s emotional state. In SZ, such perceptive abilities are blunted, and frequently associated with “negative symptoms” such as social withdrawal, flattened affect, and alogia (8,9). Patients with these negative symptoms also experience a high degree of isolation (10). Presumably, such communication impairments lead patients, by choice or by circumstance, to both *feel* isolated (i.e. lonely) and actually *be* isolated. Therefore, understanding the relationships between loneliness, desire to connect with others, social communication abilities and their causal impact on social withdrawal and isolation are particularly important.

Loneliness, independent of actual levels of social contact, plays a mediating role in physical and psychological functioning (11). Loneliness is also implicated in the development of psychosis, depression, and anxiety (12–16), and is a leading contributor to poor life satisfaction (16,17). Loneliness also exerts strong effects on cognitive functions impaired in SZ, such as executive control and attention (18,19). Importantly, loneliness and objective social isolation do not always correlate; for example, lonely individuals tend to spend just as much time with others as non-lonely people (20). Some individuals are content living solitary lives, while others feel lonely despite having many social relationships. In SZ, loneliness is frequently reported in clinical (21) and community-based studies (22), and plays a mediating role in depression in late-onset psychosis (23,24), and in adaptation to deinstitutionalization (25,26).

Whereas loneliness represents a negative emotional response to perceived isolation or a discrepancy between desired and actual social relationships (27), the "need to belong” (NTB) construct captures individual differences in the motivational desire to collaborate with others and form interpersonal bonds (28,29). It follows that NTB magnitude might influence interpersonal communication.

A prime channel through which humans communicate their social and emotional intentions is by modulating vocal intonation in speech (prosody). In healthy individuals, empathic accuracy and affective prosodic abilities positively correlate with increasing NTB ratings (30). Individuals with SZ demonstrate large effect-size deficits (Cohen’s d>1.2) in prosodic perception (31). These prosodic deficits, in turn, significantly correlate with profound pitch-perception deficits (32,33). Cues like fundamental frequency (pitch) are crucial for perceiving social intent (34,35).

Pitch perception and NTB represent two differing modulators of social communication abilities: NTB taps social-affective and a presumably stable personality trait, whereas pitch acuity represents a bottom-up sensory-processing ability necessary for prosodic processing. Pitch and prosodic acuity might influence a person’s need to connect with others or, alternatively, be the product of such needs (30,36). Regardless of the directionality of this relationship, whether individuals with SZ experience loneliness and the desire to connect with others to the same extent as healthy individuals, and whether these feelings contribute to their socio-cognitive perceptual ability, has not been studied.

In this study, we conceptualized social functioning as comprising (i) internal feelings and needs, such as desire to seek connections with others (NTB), along with self-perceptions of isolation (loneliness), and (ii) the ability to execute social interactions, such as perception of social intent through prosody, and its foundational component, pitch perception. Though avoidance of social interactions is often a feature of SZ, we hypothesized that the desire to connect with others may still be intact. Clinically, SZ patients often report feeling lonely, and Kring and colleagues found that the inner emotional lives of patients remain fairly intact (37). Consequently, we predict that SZ patients will endorse at least normal, but likely elevated levels, of loneliness. Pickett argued that individuals with a high NTB may be more inherently attentive to social cues (30). Therefore, we additionally hypothesize that prosodic abilities should positively correlate with NTB ratings. Finally, in our patient sample, we also explored the degree to which personality traits such as loneliness and NTB relate to both social communication ability and clinical symptoms such as anhedonia/asociality and avolition/apathy.

## 2. MATERIALS AND METHODS

### 2.1 Participants

Thirty-six individuals with SZ and 33 healthy controls (HC) participated. Age, gender, and parental education were comparable across groups (Table 1). Participants 1) were aged 18 to 60 years; 2) demonstrated normal hearing, according to the American Academy of Otolaryngology-Head and Neck Surgery ‘5-minute Hearing Test’; and 3) had English as their first language. Participants were recruited from a clinical participant pool at the University of Pennsylvania, and were solicited through community postings and outpatient clinics. Patients who participated 1) met criteria for SZ (n=31), schizophreniform (n=3), or schizoaffective disorder-depressive type (n=2) as determined by the Structure Clinical Interview for DSM Disorders (SCID) administered by a trained clinician in the preceding year; 2) were deemed clinically stable by their consultant; and 3) had been on a stable medication regime unchanged over the past three months. According to the SCID, controls showed no evidence of past or present mental illness, 1) were not taking psychoactive medication, and 2) did not have a first-degree relative with psychosis. The University of Pennsylvania Institutional Review Board approved study procedures. Written informed consent was obtained from all participants.

### 2.2 Procedures

For each participant, the order of scales and experimental tasks was randomized.

#### 2.2.1 Scales

Participants completed scales alone. Loneliness was assessed using the UCLA Loneliness Scale – Revised, a 20 item questionnaire designed to measure perceived isolation (38), and comprising three subscales: isolation (social dissatisfaction at the individual level) and relational- and collective-connectedness (satisfaction with social self at the relational and collective levels) (39). Necessary items were reverse-coded to calculate a total score that reflects the overall level of loneliness, up to a maximum score of 80, with a published mean score for healthy young adults 40.0 (38). Desire for social connectedness (NTB) was assessed using the Need to Belong Scale (28,40,41), a 10 item scale (maximum score of 50; published mean score for healthy young adults 32.8; 40) that attempts to quantify an individual’s need for positive social contacts and acceptance or aversion of rejection,. Two participants with SZ were excluded at this stage because of lack of consistency of responses between reversed and non-reversed items on the Loneliness Scale. Six individuals (5 SZ) omitted answering one question on the Loneliness Scale. These missing items were assigned the average value of responses for the relevant dimension (39).

#### 2.2.2 Pitch Perception

Pitch perception was assessed using the measures of Leitman et al. (42). Participants were presented with pairs of 100 ms tones with 500 ms inter-tone intervals. To avoid learning effects, three different base tones (500, 1000, and 2000 Hz) were incorporated. Of the 94 stimuli pairs, half were identical, and half differed in frequency by 1%, 1.5%, 2.5%, 5%, 10%, 20%, or 50% to their counterpart. Participants were asked to indicate whether the pitch was the same or different. One participant responded with "same” for every item, and therefore their data from this task were excluded.

#### 2.2.3 Prosody task

We assessed prosody perception using the Auditory Emotion Recognition task, comprising a subset of stimuli from Juslin and Laukka, which utilizes recordings of British English speakers vocalizing neutral sentences (e.g. "it is 11 o’clock”) with angry, fearful, happy, sad, or neutral intonation (35). Each emotion was presented six times. Because large variation in pitch most classically predicts happiness, this emotion was presented nine times, to assist in detecting the relationship with pitch perception. From these forced-choice response options, participants were asked to choose which emotion was conveyed by intonation.

#### 2.2.4 Clinical symptom measures

Clinical symptoms were measured using the SANS (Scale for the Assessment of Negative Symptoms; 43) and SAPS (Scale for the Assessment of Positive Symptoms; 44).

### 2.3. Data Analysis

Multivariate analyses of between-group endorsements of loneliness, NTB, and performance on the pitch and prosody tasks were conducted. Effect sizes (Cohen’s d) were calculated and interpreted according to Cohen’s guidelines (45). Analysis of sample distribution densities/shape differences was conducted using Q-Q analysis, with a Shapiro-Wilk assessment to test for normality. Correlations between measures were calculated using Spearman correlations. All statistical tests were conducted in R (http://www.r-project.org), with a <0.05 two-tailed α criterion.

## 3. RESULTS

### 3.1 Scales

SZ participants reported feeling lonelier than HC on the UCLA Loneliness Scale, (t_57_ = - 8.37, p<0.01; Cohen’s d=1.43) (Figure 1A). However, SZ participants scored comparably to HC on the Need to Belong Scale (NTB) (t_62_ = -0.61, p>0.54; d=0.15) (Figure 1B). Q-Q plot analysis indicated similar rating distributions for both groups on the NTB. A Shapiro-Wilk test further demonstrated normality of distribution of NTB data across groups (w = 0.98, p>0.36). However, relative to controls, SZ patients’ loneliness ratings were less positively skewed, less leptokurtic, and less restricted in their distribution range. Here, a Shapiro-Wilk test showed non-normal data (w = 0.96, p<0.04). Loneliness and NTB scores did not correlate within the SZ (r_s_ = -0.17, p>0.34) or HC groups (r_s_ = 0.00, p>0.98). (Figure 1C). Examination of loneliness endorsements along the UCLA scale’s three dimensions — isolation (t_58_ = -7.08, p<0.01), relational connectedness (t_53_ = -7.46, p<0.01), and collective connectedness (t_55_ = -4.69, p<0.01) — indicated large effect-size increases in SZ endorsements for all dimensions (all d> 1.0) (Figure 1D).

**Figure 1.**
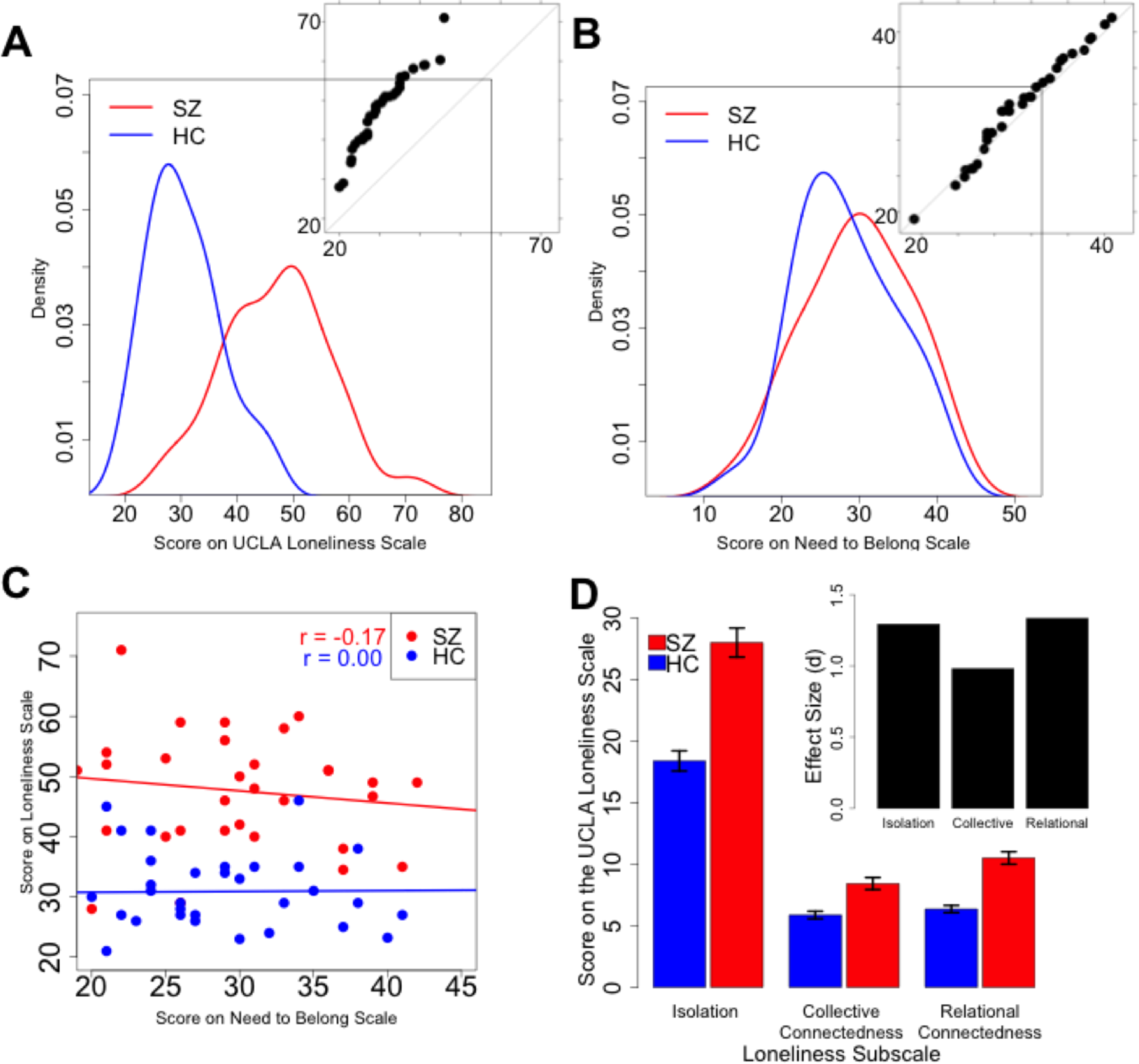
Loneliness and Need to Belong Scores. A) Density plots and qqplot (inset) show higher loneliness scores in SZ participants compared to HC participants. B) Density plots and qqplot show similar NTB ratings between groups. C) NTB and loneliness scores showed no correlations in either group. D) UCLA Loneliness Scale endorsements according to its three conceptual dimensions: general isolation, relational connectedness, and collective connectedness. Note that reverse coding of items results in higher scores in relational and collective connectedness, indicating a lack thereof. Error bars represent the standard error. Inset: Effect sizes of differences between SZ and HC groups.

### 3.2 Prosody and pitch perception

SZ participants were less able to correctly identify emotions from prosody compared to HC (t_61_ = 3.11, p<0.01; d=0.73) (**Table 1**). Prosody task scores correlated positively with NTB scores only in SZ (r_s_=0.45, p<0.02) (Figure 2A). Correlations between prosody and loneliness were not significant (Figure 2B). SZ participants correctly identified fewer pitch task trials compared to HC (t_59_ = 4.91, p<0.01; d=1.04) (**Table 1**). Pitch perception was related to prosody perception within the SZ (r_s_ = 0.41, p<0.02) and HC groups (r_s_ = 0.49, p<0.01). No other correlations between task performance and scale scores were significant (p >0.17).

**Figure 2.**
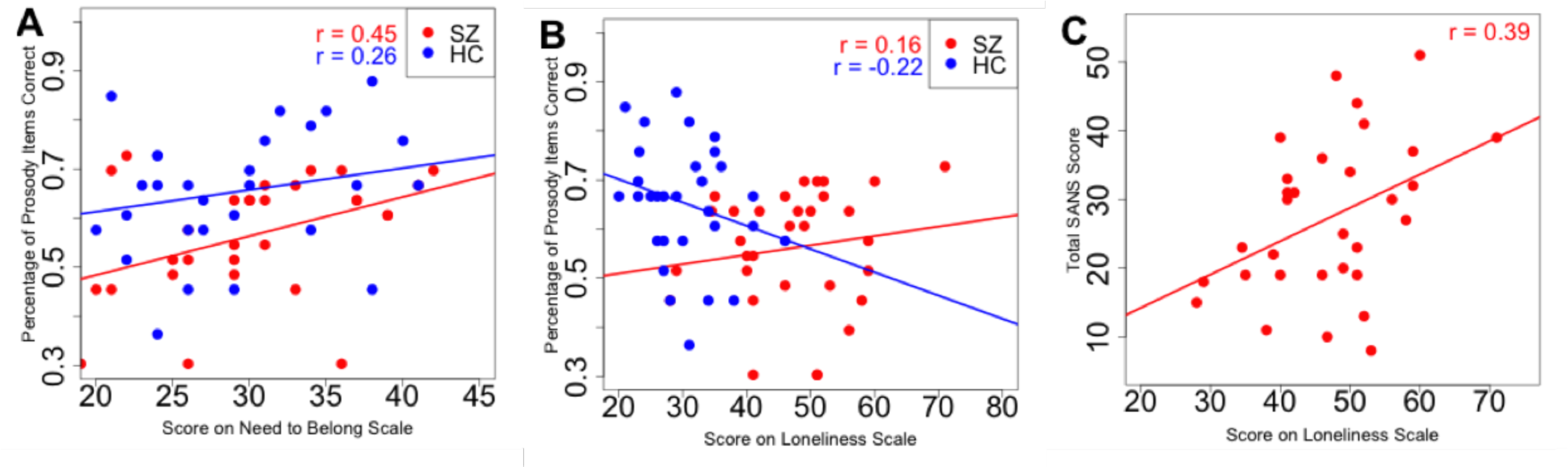
Relationships of Loneliness and Need to Belong to behavioral performance and clinical symptoms. A) NTB scores and prosody task performance positively correlated in SZ. This correlation was not statistically significant in controls. B) Correlations between loneliness scores and prosody task performance were not significantly correlated in either group. C) Loneliness scores correlated with overall negative symptomology as determined by the SANS.

### 3.3 Relationships with clinical and demographic measures

Within SZ, endorsements of loneliness (r_33_ = 0.39, p<0.03), but not NTB, correlated positively with clinical ratings of Negative Symptoms (SANS) (Figure 2C). Scores on the anhedonia/asociality subscale also correlated positively with endorsements of loneliness (r_33_ =0.53, p<0.01). No other correlations between task performance, scale scores, and symptom sub-scales were observed (all p’s >0.05). Antipsychotic medication dosage (CPZ equivalency) did not correlate with any of the tasks, scales, or clinical indices (all p’s >0.13) (46).

## 4. DISCUSSION

Consistent with clinical reports, patients with SZ endorsed higher levels of loneliness (d=1.8) compared to HC. However, ratings of desire for social connectedness (NTB) among SZ patients were normally distributed and equivalent to those of HC (d=0.2), indicating similar degrees of heterogeneity in HC and SZ groups in the desire to connect with others. As previously demonstrated, pitch acuity and prosody perception were correlated, confirming the importance of basic sensory processing in recognizing prosodic emotions. Scatterplots indicated loneliness and desire for social connectedness to be independent factors, and that socio-cognitive perceptual ability (through prosody) correlated with increased desire for social connections.

Our finding of normally distributed desire for social connectedness is consistent with Trémeau et al., who observed normal levels of "social” but not “situational” motivation in SZ using the Motivation and Energy Inventory (47). At face value, intact desire for social connectedness may seem to contradict classic conceptions of social withdrawal and anhedonia. However, Trémeau, as well as Kring and Elis, found little evidence to suggest a patient’s experiential emotional world precludes him/her from feeling the desire to connect with others (48,49). Furthermore, normal desire for social connectedness is consistent with arguments that fundamental human motivations such as desire to belong are adaptive regardless of a person’s current state of belonging (28,40).

Our findings of increased loneliness are in line with prior clinical observations and self-report studies (21,22,50). Prior research indicated that individuals with SZ report high levels of loneliness, both in the early onset of the disease (51) and after a period of stability (52,53). Though these feelings of loneliness may reflect higher levels of objective isolation experienced by SZ patients relative to healthy individuals, prior studies in non-clinical populations indicate a strong disjunction between perceived isolation (loneliness) and actual isolation in terms of independent effects on social and functional satisfaction and mental and physical health (18).

To better understand how these concepts are related, an objective measure of isolation is required. While it has been speculated that loneliness and desire for social connectedness are closely related (54), recent work in healthy individuals indicated only 7.8% of shared variance between the UCLA Loneliness and NTB scales (41). Both lonely and non-lonely individuals may have the same number of social interactions, suggesting that subjective feelings of isolation, rather than actual number of social interactions, are the cause of the detrimental social and emotional changes (55).

Baumeister and Leary and Pickett et al., argue that scales like the NTB scale index the degree to which an individual feels this desire, and that individuals with higher desire for social connections develop higher levels of sensitivity to social cues like prosody (28,30). Our prior studies suggest that the causal arrow may actually point in the opposite direction, such that impairments in pitch perception may lead to functional socio-cognitive perception impairments (31,56). Moreover, recent studies indicate that perception of social cues from prosody is impaired even before formal diagnosis is reached (57,58), and before long-term deterioration in social function has become ingrained (59). These support a clinical trajectory in which sensory-based disturbance upwardly generalizes into social communication impairment, whose sequelae include social withdrawal isolation and loneliness. Finally, research indicates psychosis can both lead to, and be produced by, isolation and loneliness (60,61). In SZ, processing dysfunction beginning at early stages could influence perception of social cues from prosody, consequently generating misinterpretation. The response to impaired socio-cognitive perception could likely be social withdrawal, resulting in first objective isolation and, consequently, loneliness. A cycle might then ensue, with self-imposed isolation and loneliness further exacerbating symptoms, and degrading social skills further. While it could be assumed patients with severe negative symptoms might not be subjectively distressed by the impact of social withdrawal, recent evidence suggests negative symptoms can develop as a reaction to self-defeatist beliefs such as loneliness (62,63). In particular, loneliness displays a close relationship with anhedonia, which might reflect the role of amotivation and effort in the latter (64).

The cross-sectional nature of our study precludes conclusions as to causal direction, but this could be established with a longitudinal design. In any case, understanding the relationship between desire for social connectedness, feelings of loneliness, actual isolation, and socio-cognitive perception is of great practical importance, and we did observe a relationship between severity of negative symptoms and perceived loneliness. That desire to belong did not correlate with negative symptoms is perhaps not surprising, given that patients did not demonstrate impaired desire for social connectedness.

Modeling how these factors interact across psychiatric illnesses in which social dysfunction presents may provide further clues as to how behavioral traits combine with sensory and/or cognitive processing impairment to produce social communication dysfunction, as well as clues for remediation strategies. For example, based on our data and that from other research on normal and autistic populations, we present theoretical models for remediating socio-cognitive perceptual impairment in Figure 3. We contrast our model of social function in SZ with a possible model for another disorder with well-known social impairments, namely autism. For a full discussion, see figure caption.

**Figure 3.**
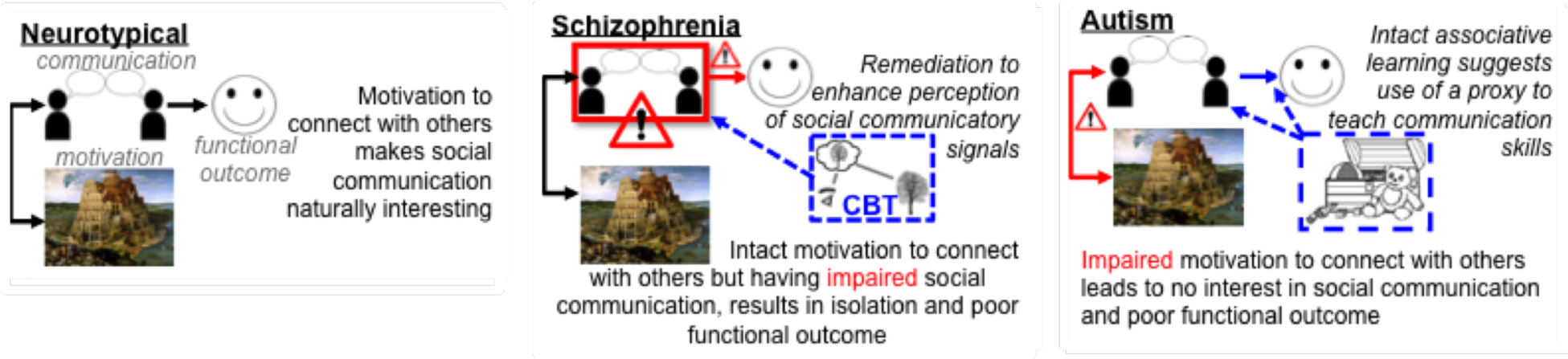
Theoretical models for remediating socio-cognitive perceptual impairment in neuropsychiatric disorders. Neurotypical individuals are naturally motivated to find social interactions reinforcing. In SZ, our data suggest that not only is the desire for social connections intact, but there is also an association between this desire for social closeness and the ability to correctly interpret prosody. Thus, if socio-cognitive perceptual impairments exist, but desire for social connectedness remains, interventions to improve perceptual deficits could be an effective means to improve social functioning. The effectiveness of social skills training appears promising for people with SZ (65,66), as might behavioral training as has been used successfully in other populations to improve pitch acuity (e.g. musicians) (67). Additionally, the success of interventions to reduce loneliness in affected populations (68) also holds promise for improving social functioning in SZ, in addition to enhancing social support through regular contacts or companionship. On the other hand, different disorders may follow different pathways to these deficits despite showing similar functional outcomes. Social functioning deficits in Autism Spectrum Disorders (69,70) could result from impaired social motivation (71). Therefore, children with autism may not automatically pay attention to social cues or find them rewarding. In this autism model, in contrast to the model for SZ, treatment should instead focus on reinforcing socio-cognitive perceptual abilities by pairing social interactions with a desired reinforcer, thus mimicking increased desire for social connectedness.

This study, though preliminary in nature, provides a path for future research. Future studies should incorporate more extensive measurements of social skills. To clarify how the need to connect with others affects our functional abilities to socialize and perceive social intent, new tools and paradigms to measure the desire for social connection and loneliness are needed. Scales such as the UCLA Loneliness and NTB Scales, while useful, rely on introspection and self-report. True characterization of the relationship between desire for social connectedness and socio-cognitive perceptual ability may only come through developing objective tasks and paradigms that can manipulate the desire to connect. Such tasks may help delineate the nature of desire for social connectedness, and ascertain causal relationships between desire for social connections, loneliness, and socio-cognitive perceptual abilities that cannot be inferred through correlations alone.

In conclusion, the data presented here suggest individuals with SZ have intact desire for social connectedness, and we speculate that feelings of loneliness and actual isolation may partly result from impaired socio-cognitive perception. This conclusion could illuminate different models of sociality within different disorders, for which more targeted treatments could be developed.

